# Genome-scale community metabolic modeling of maize root-associated microbiota shows that root exudates stimulate diverse metabolic interactions

**DOI:** 10.64898/2026.05.13.724839

**Authors:** Ashley E. Beck, Henry Phillip, Anna-Katharina Garrell, Manuel Kleiner

**Affiliations:** Department of Biological and Environmental Sciences, Carroll College, Helena, Montana, USA; Department of Plant and Microbial Biology, North Carolina State University, Raleigh, North Carolina, USA

## Abstract

Microbes play a vital role in plant development, health, and resilience, yet relatively little is known about the specific metabolic mechanisms driving interactions in these host-associated communities. Systems biology models enable a computational approach to understanding metabolic interactions, which can be difficult to pinpoint experimentally; however, these methods cannot yet accommodate the large number of species in natural communities. Synthetic communities (SynComs) provide a more tractable alternative to explore targeted interactions. Here, we investigated metabolite exchange in a seven-member maize root-associated SynCom, specifically accounting for plant host context by designing a customized exudate medium. We constructed metabolic models for each bacterial species and curated them with *in vitro* phenotyping data to reflect experimentally based carbon uptake potential. Flux balance analysis of individual species demonstrated that integrating phenotype data and changing medium type had substantial impacts on predicted growth rates, which in turn shaped potential interspecies interactions. *In silico* community growth optimization of the seven-member community model showed that the exudate medium supported a more diverse community composition compared to minimal medium, with predictions of community member abundance closely aligned to literature-derived experimental results. Predicted metabolite exchange in the root exudate environment showed *Enterobacter ludwigii* as a community hub, and cross-feeding of indole suggested a potential effect of bacterial community interactions on the plant host. Our *in silico* findings indicate the host plays an important role in structuring microbial interactions and cross-feeding at the metabolic level, underscoring the importance of considering environmental context from both theoretical and experimental perspectives.

**IMPORTANCE:** True understanding of a system is marked by the ability to predict its behavior. The complexity of natural host-microbe systems represents a frontier of knowledge that scientists are working to understand, and elucidating principles of interactions within multi-partite microbial communities remains a challenge in microbial ecology. Synthetic communities provide a tractable starting point for investigating interaction mechanisms, and computational approaches complement laboratory experiments by systematically evaluating multiple possibilities for metabolic pathway processing, thereby allowing us to comprehensively study the interconnected metabolic networks of host-associated microbiota. The model we developed for the seven-member maize root-associated bacterial community presents a step toward predicting plant-microbe behavior, providing hypotheses for future experimental testing and serving as a template for expanding model complexity to more members and other systems.

## INTRODUCTION

Plant-microbe interactions play key roles in plant development, health, and resilience to biotic and abiotic stresses (1–4). Despite the essential role of the plant microbiota, the mechanisms by which microbial community members interact with each other to colonize plant roots and assemble into diverse and stable communities remain unclear (5). Resource competition and metabolic exchange are known to be important determinants of microbial community interactions (6, 7), but the manner in which the plant host influences these interactions is unknown. A recent study modeling microbe-microbe interactions in the *Arabidopsis* phyllosphere demonstrated the impact of niche overlap on competitive interactions (8), underscoring the need to understand the metabolic capabilities of individual members to probe community interactions. While *Arabidopsis* is an important model plant, expanding research to elucidate drivers of microbial community stability on or within agricultural plant hosts is key to establishing improved agricultural practices and enhancing conservation and sustainability efforts.

*Zea mays* (maize) is a globally important cereal crop and is becoming a model for investigating plant-microbe interactions (9–14). Studies have characterized microbiota composition in a variety of tissues, observing differences in response to soil characteristics, plant host genotype, stage of plant development, and agricultural practices (15). While microbes colonize all plant tissues, the root-associated microbiota is of particular interest given its role in nutrient acquisition (16). Previous work has generated a synthetic community (SynCom) representative of the maize root microbial community. This community consists of seven bacterial species isolated from *Zea mays* cv. Sugar Buns (12) and provides a reduced complexity, tractable experimental system for studying microbial interactions in the root system. In their study, Niu et al. (12) demonstrated that the removal of a single species – *Enterobacter ludwigii* – triggered community collapse, which led them to speculate that *E. ludwigii* may represent a keystone species that has a critical influence on community structure. Furthermore, a recent study using this SynCom (14) identified unique metabolic niches for each species via exometabolomic profiling of cultures supplemented with maize root extract. Cross-feeding experiments in this study also identified the potential for interactions based on prototroph-auxotroph relationships.

Stoichiometric metabolic models encapsulate the metabolic potential of an organism based on genome sequence and provide a systematized approach to analyzing metabolite conversion capabilities (17), which can be extended to multiple species to examine and predict interactions in a community. The main challenges to address in this area are effective integration of individual models into a community model and appropriate representation of the experimental context, such as a plant host (18). With advances in both genomics and computational capacity, a number of studies have explored modeling microbial consortia for bioprocess or nutrient conversion involved in geochemical cycles, bioremediation, and wastewater treatment (19–22), along with host-related studies focusing primarily on human gut microbiota (23). While a few studies have directed modeling efforts toward plant microbiota (*e.g.*, Schäfer et al. (8)), explicit consideration of the host environment, as well as ecological concepts such as resource utilization principles (18), are still needed. Such an approach will enhance the utility and relevance of community metabolic models. Principles and mechanisms postulated with SynComs, via both computational and laboratory approaches, can then be tested or scaled up under field-relevant conditions to validate hypotheses.

In our study, we constructed and analyzed genome-scale metabolic models for the Niu et al. (12) seven-member maize root SynCom. We examined the impact of incorporating experimental phenotyping data on model predictive capability for each of the seven community members individually, as well as the influence of growth medium environment by comparing our custom-designed root exudate medium with a minimal laboratory cultivation medium (24). Following these analyses, we combined individual models to create a compartmentalized community-scale model and assessed the impact of medium environment on predicted community structure. We found that proper consideration of environmental context through medium design produces predictions that better align with experimental measurements of community abundance, and the predicted exchanges of amino acids, nucleosides, and indole provide hypotheses for future experimental testing of community interactions.

## RESULTS

### Designing a contextually relevant simulation environment

Accurately representing the maize root environment in model simulations was central to the study design to provide real-world relevance. In lieu of adding a multicellular plant to the system of models (*e.g.*, (25)), which would substantially increase computational and analytical complexity, we designed a medium containing a suite of maize root exudate compounds (based on Hao et al. (26)) to emulate the root environmental context; for comparison, we used a previously designed minimal medium that was developed specifically to enable growth of each of the SynCom species in the laboratory (24). While it was not feasible to include every possible metabolite in the exudate medium, we augmented the substrates to realistically reflect rhizosphere complexity and variety, including 11 sugars, three alcohols, 10 organic acids, nine amino acids, and two pyrimidine nucleosides (**Table 1**); only metabolites with corresponding BiGG standardized identifiers were used to maintain compatibility with CarveMe model construction software (27). We classified compounds as either “primary” or “secondary” substrates based on their likely contribution to carbon catabolism: sugars, alcohols, and organic acids are likely the main sources of carbon and were considered primary substrates, while amino acids, nucleosides, and vitamins were considered secondary. Substrate concentrations were not directly factored into the medium, as stoichiometric steady-state modeling algorithms do not account for the dynamic nature of substrates in the environment; however, substrate uptake constraints were set to reflect a higher relative availability of primary compared to secondary substrates in some simulation scenarios, as described in the next section. A complete composition list for both media (referred to hereafter as minimal and exudate media) is found in Table S1-2 of Supplemental File S1.

**Table 1.** Comparison of medium components between the standard minimal medium for the SynCom (24) and our customized representation of a natural maize root environment (based on secreted compounds identified in Hao et al. (26)). Compounds unique to each medium are highlighted in the “Minimal Medium Only” and “Exudate Medium Only” columns while shared compounds are shown in the center column. Glucose and malate serve as the central carbon sources for the minimal medium, whereas the exudate medium was designed to encompass a variety of other representative sugars, alcohols, acids, and nucleosides. Sugars, alcohols, and organic acids were designated as primary substrates (1°, main contributors to carbon catabolism), while amino acids, nucleosides, and vitamins were designated as secondary substrates (2°). Nitrogen (ammonium, nitrate), phosphate, and sulfate were also included in both media, along with key metals and ions (full composition list available in Table S1-2 of Supplemental File S1).

| Substrate Type | Minimal Medium Only | Both Minimal and Exudate Media | Exudate Medium Only |
| --- | --- | --- | --- |
| Sugars (1°) | - | glucose | arabinose, erythrose, fructose, galactose, isomaltose, lactose, maltose, melibiose, sucrose, xylose, |
| Alcohols (1°) | - | - | glycerol, inositol, ribitol |
| Organic acids (1°) | - | malate | cis-aconitate, citrate, benzoate, fumarate, lactate, malonate, oxaloacetate, succinate, tartaric acid |
| Amino acids (2°) | C, K, M, P | I, L, N, Q, R, S | A, D, E, G, GABA, H, T, V, Y |
| Nucleosides (2°) | - | - | cytidine, thymidine |
| Vitamins (2°) | B2, B3, B5, B6, B7, B12 | - | - |

### *In vitro* metabolic phenotyping data and medium type alter predicted growth rate in accordance with carbon uptake capacity

The metabolic capabilities of bacterial species are commonly gathered from literature, or generated in the laboratory if possible, for validation and curation of metabolic models. The Biolog microplate method is widely used to analyze metabolic capabilities of microorganisms (*e.g.*, (28, 29)). We used the Gen III microplate, designed for aerobic bacteria, to assess carbon metabolism potential across a range of compounds for each of the seven root-associated bacterial species from the Niu et al. (12) SynCom: *Brucella pituitosa*, *Chryseobacterium indologenes*, *Curtobacterium pusillum*, *Enterobacter ludwigii*, *Herbaspirillum robiniae*, *Pseudomonas putida*, and *Stenotrophomonas maltophilia* (**Table 2**). The seven species showed a variable range of substrate metabolism capacities, with *E. ludwigii* being the most versatile and *S. maltophilia* the least. We used this data in conjunction with CarveMe software (27) to curate a genome-scale metabolic model for each species.

**Table 2.** Substrate metabolism capability for each bacterial species as experimentally measured with a Biolog Gen III microplate (n = 3 for each species). While some of the compounds could also serve as nitrogen sources, the Biolog assay tests substrate efficacy as sole carbon and energy sources. Blue (+) indicates growth and therefore uptake and metabolism of the compound; pink (−) indicates no growth on the carbon source. Only substrates relevant to the minimal and exudate media are shown here; Supplemental File S2 includes the complete data set. The last row denotes the total number of positive results for each species from the presented subset.

| Substrate | *B. pituitosa* | *C. indologenes* | *C. pusillum* | *E. ludwigii* | *H. robiniae* | *P. putida* | *S. maltophilia* |
| --- | --- | --- | --- | --- | --- | --- | --- |
| Maltose | + | + | + | + | - | - | + |
| Sucrose | - | - | + | + | - | - | - |
| Lactose | - | - | + | + | - | - | - |
| Melibiose | - | + | + | + | - | - | - |
| Glucose | + | + | + | + | + | + | + |
| Fructose | + | + | + | + | - | + | - |
| Galactose | + | + | + | + | + | - | + |
| Inositol | + | - | + | + | - | - | - |
| Glycerol | + | + | + | + | + | + | - |
| Proline | + | + | + | + | + | - | + |
| Alanine | + | + | + | + | + | + | + |
| Arginine | + | + | + | + | - | + | - |
| Aspartate | + | + | + | + | + | + | + |
| Glutamate | + | + | + | + | + | + | - |
| Histidine | + | + | - | + | - | + | - |
| Serine | + | + | + | + | - | + | + |
| Lactate | + | - | - | + | + | + | + |
| Citrate | - | + | + | + | + | + | + |
| Oxaloacetate | + | - | - | - | + | + | - |
| Malate | + | - | - | + | + | + | + |
| Succinate | + | - | - | + | + | + | + |
| GABA | + | - | - | - | + | + | - |
| Total positives | 18 | 14 | 16 | 20 | 13 | 15 | 11 |

We constructed genome-scale metabolic models for each bacterial species in CarveMe (27) with gapfilling on either the minimal or exudate medium (**Table 1**). For each medium, two separate models were constructed: either (1) incorporating Biolog phenotype data as soft constraints, which prioritizes alignment of specified uptake reactions with corresponding experimental data (27), or (2) excluding the integration of Biolog data. Taken together, four different model versions were constructed for each of the seven species (illustrated in **Figure 1**); these parallel models allowed for distinguishing how simulation output was affected by (1) integration of phenotyping data and (2) the medium selection. Basic model statistics for each model version including numbers of reactions, metabolites, and genes are included in Table S3 of Supplemental File S1.

**Figure 1.**
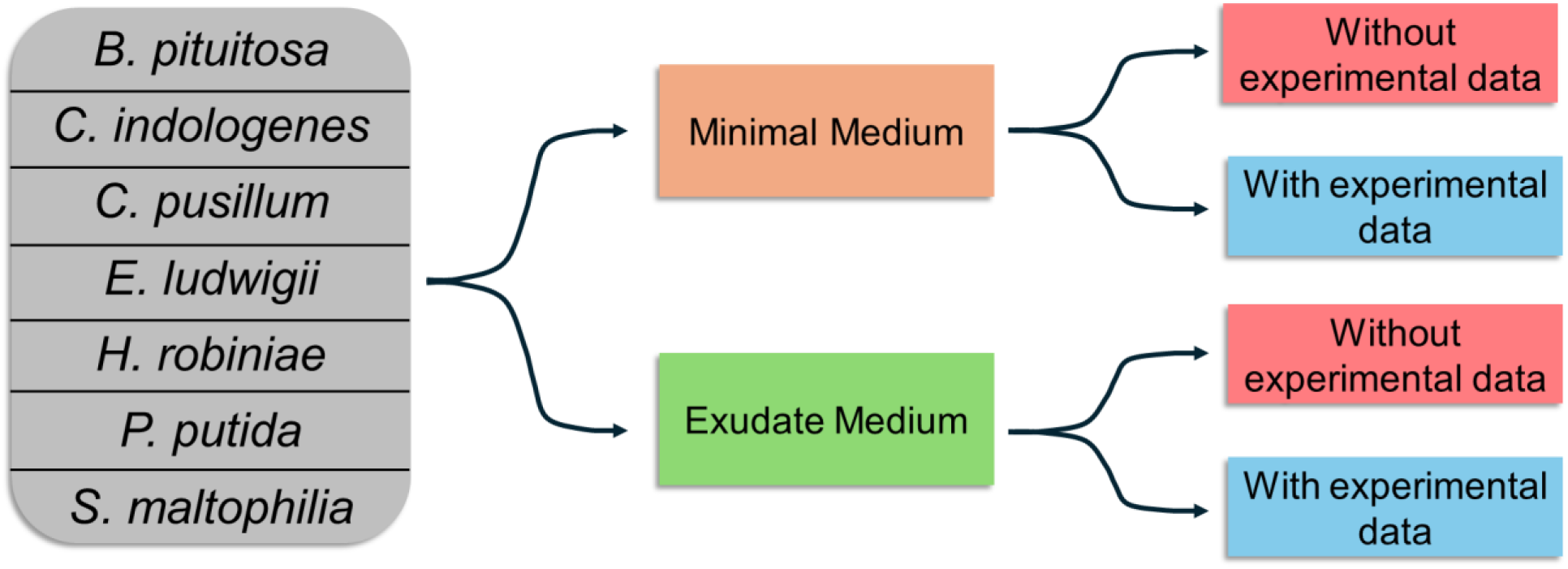
Illustration of the parallel model versions constructed for each bacterial species in this work and explanation of main simulation parameters. For each of the seven species listed, models were constructed separately in CarveMe with the appropriate Gram-positive or Gram-negative template and genome sequence, using either the minimal or exudate medium as the gapfill environment. In each case, models were constructed either with or without the incorporation of Biolog metabolism data as soft constraints. This approach generated 28 models in total (four different versions for each species), to which flux balance analysis (FBA) was then applied. The substrate uptake constraints applied were adjusted according to the intent of the simulation. (1) To assess the effect of incorporating phenotyping data, the uptake was set to 10 mmol·gCDW^−1^·h^−1^ (CDW, cell dry weight) for each substrate to avoid potential bias of nutrient limitation. (2) To evaluate differences between media, a total uptake of 10 mmol·gCDW^−1^·h^−1^ was partitioned with equal distribution across the primary substrates (sugars, alcohols, and organic acids), and similarly, a total uptake of 0.25 mmol·gCDW^−1^·h^−1^ was partitioned equally across the secondary substrates (amino acids, nucleosides, and vitamins). Simulations optimized biomass production as the objective function, *i.e.*, the metabolic goal of the cell.

Comparing model versions contextualized with or without Biolog data for each medium type revealed the influence of experimentally determined substrate uptake capacity on model predictions. We performed flux balance analysis (FBA) simulations to optimize for growth (**Figure 2**), using a default uptake rate of −10 mmol·gCDW^−1^·h^−1^ (CDW, cell dry weight) for each substrate to provide a direct comparison of predicted growth rates without imposing severe nutrient limitations. Corresponding simulations including and excluding Biolog data for each species were compared by normalizing both growth rates to that of the data-excluded simulation 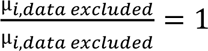 and 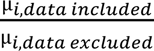 for each medium type, where µ is the predicted growth rate and 𝑖 represents the species). This scaling approach allowed observation of overall trends and evaluation of the differential impact across species and media. Some species showed similar growth rate predictions with and without phenotyping data, while others exhibited substantial differences; it should be noted that each simulation resulted in a slightly different maximal growth rate. Similarities were more prevalent in the minimal medium (**Figure 2A**), which contains fewer components than the exudate medium, while greater variation was evident under the exudate medium condition (**Figure 2B**). The average and median percent differences between predicted growth rates including and excluding Biolog data were 18% and 15%, respectively, for the exudate medium, compared to an average of 9% and a median of 1% for the minimal medium.

**Figure 2.**
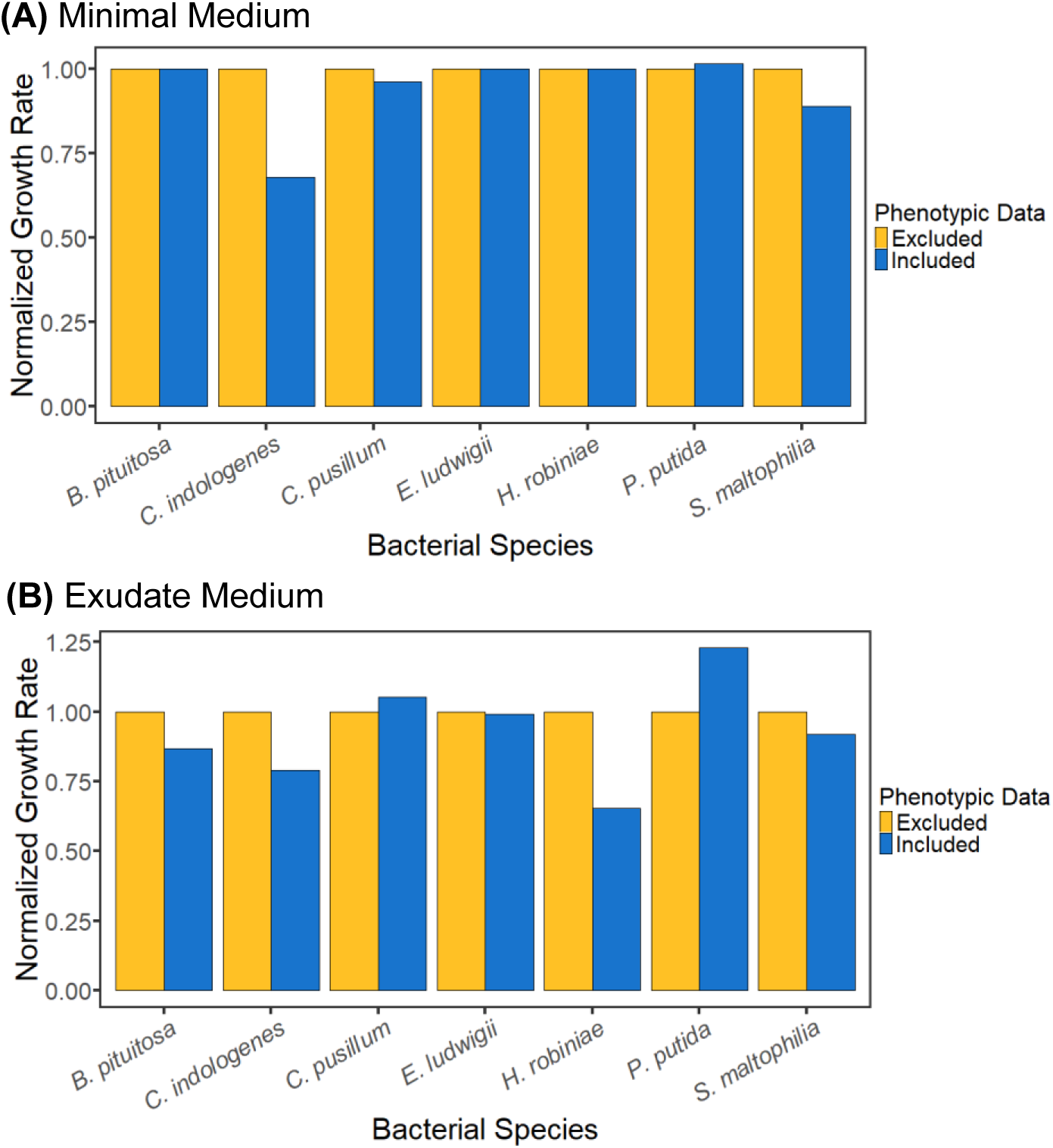
Effect of integrating Biolog phenotyping data on simulated growth rates in **(A)** minimal medium and **(B)** exudate medium conditions for the seven individual metabolic models, representing monoculture growth. Biolog data regarding substrate uptake and metabolism were integrated as soft constraints in one set of CarveMe models (blue series, “Included”) and not incorporated in a parallel set (yellow series, “Excluded”). FBA was used to optimize each model for maximal growth. Simulation results are plotted as the normalized growth rate comparing the two model versions, scaling by the growth rate obtained without data integration for each pair of simulations; this scaling establishes a frame of reference for assessing the magnitude of impact across species and between media. Maximum uptake for each substrate was set to −10 mmol·gCDW^−1^·h^−1^ to avoid unintended bias from nutrient limitation.

Incorporating these experimentally observed capabilities can have an effect on optimal growth rate as observed in **Figure 2** (see (30) for another example), but the direction of the effect cannot be predicted *a priori* without knowledge of the physiological characteristics of the species. In both medium conditions, predicted growth rates were higher when Biolog data were not incorporated for five of the seven species. These differences in predicted growth rates were due to the models incorrectly predicting substrate consumption, likely due to discrepancies or inconsistencies in transporter identification (31). For example, the two primary carbon sources in the minimal medium are glucose and malate, and experimentally, *C. pusillum* was unable to consume malate but could take up leucine and serine. Without including this experimental data, however, the optimal flux pathway predicted maximum malate uptake and no leucine or serine consumption, contributing to a 4% higher growth rate when experimental data were excluded (**Figure 2A**). As another example, *H. robiniae* showed a substantial increase in predicted growth rate when phenotyping data were excluded from the exudate medium; simulation results without data integration predicted uptake of fructose, maltose, and sucrose, which *H. robiniae* was not experimentally capable of metabolizing and were disallowed in the simulation incorporating experimental data. In other cases, including Biolog data identified transporter or pathway gaps in the genome, resulting in a higher growth rate when experimental data were incorporated (*e.g.*, as for *P. putida* in both minimal and exudate media).

In light of the influence of integrating phenotyping data, only models incorporating Biolog data were used for subsequent simulations. To examine the influence of medium type, simulations were performed for each bacterial species separately under both minimal and exudate media conditions. The total uptake of major substrates (sugars, alcohols, and organic acids) in each medium was set to −10 mmol·gCDW^−1^·h^−1^, and the total uptake of secondary compounds (amino acids, nucleosides, and vitamins) was set to −0.25 mmol·gCDW^−1^·h^−1^. These constraints were applied to provide a more equitable scale for comparing model performance between the two media. To exemplify, in the minimal medium, the maximum uptake rate of −10 mmol·gCDW^−1^·h^−1^ was divided only between glucose and malate (−5 mmol·gCDW^−1^·h^−1^ each), whereas the exudate medium contained 24 possible primary substrates (sugars, alcohols, and organic acids). Therefore, each species was constrained to a smaller maximum uptake per substrate (−10 mmol total / 24 substrates = −0.42 mmol·gCDW^−1^·h^−1^). Otherwise, if we had set the same uptake rates for individual substrates, the total substrate availability between the two media would have differed substantially due to differing numbers of substrates.

Predicted growth rates differed between minimal and exudate media for each species (**Figure 3**) with differences ranging from 2% for *E. ludwigii* to 118% for *H. robiniae*, reflecting differences caused by the variety and amount of substrates available, along with species-specific differences in ability to metabolize available substrates. For example, *H. robiniae* was predicted to metabolize 10 additional primary substrates included in the exudate medium compared to 15 for *E. ludwigii*, contributing to the larger difference between predicted growth rates for these species. For most species, faster growth was predicted under minimal medium conditions; *C. pusillum* was the exception, with slightly faster growth predicted in the exudate medium due to its inability to consume malate (one of the two major carbon substrates available in the minimal medium). For each species, faster predicted growth rate correlated with higher total carbon substrate uptake rate (Figure S1-2 in Supplemental File S1), indicating that the advantage of one medium environment over the other is tied to each microbe’s carbon and energy uptake capacity: a larger amount of a smaller number of substrates is more efficient for growth than access to a wider array of substrates in smaller quantities. It is important to note that these results represent pure culture scenarios; the efficiency of substrate uptake strategies will likely vary when microbes coexist in community due to competition for resources, which will be explored in the community modeling section below.

**Figure 3.**
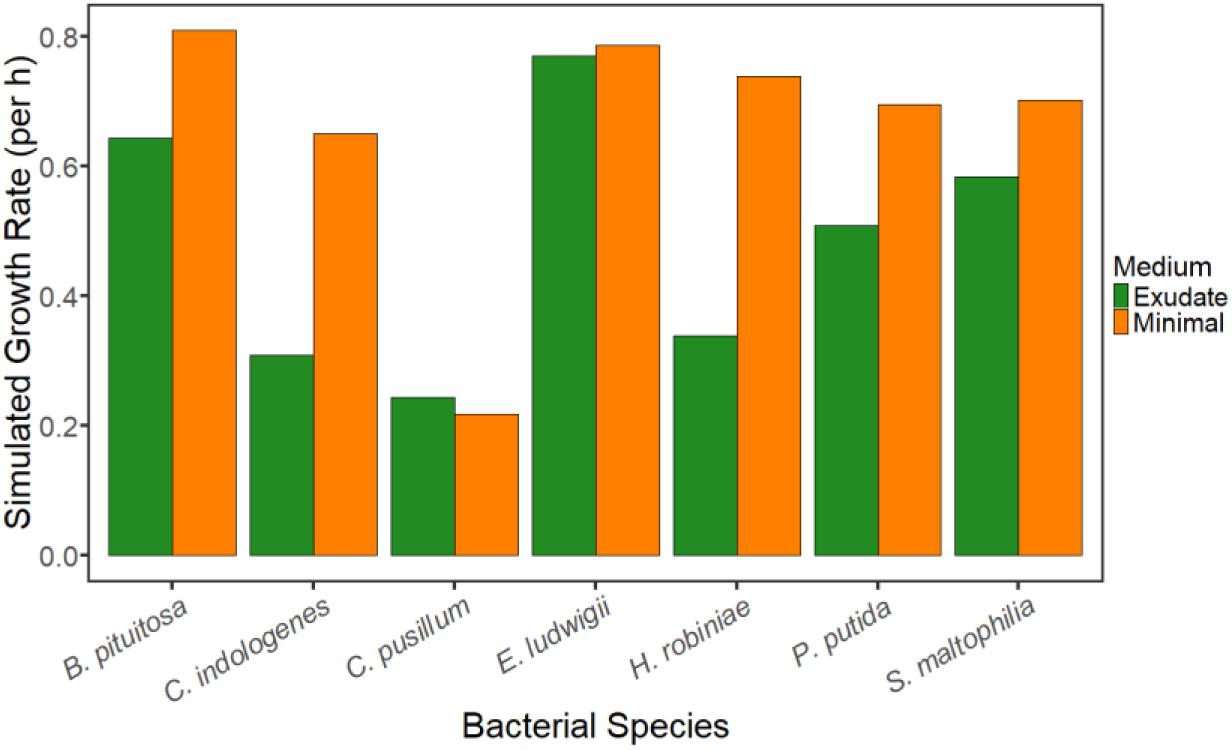
Effect of medium type on simulated growth rates, comparing the minimal and exudate media (orange and green series, respectively), with models incorporating Biolog data. Simulation results represent the growth rate-maximized objective values obtained with FBA. The total combined uptake of major carbon sources (sugars, alcohols, and organic acids) was set to −10 mmol·gCDW^−1^·h^−1^, and the total combined uptake of secondary substrates (amino acids, nucleosides, and vitamins) was set to −0.25 mmol·gCDW^−1^·h^−1^, to provide a comparable scale for assessing model performance between the two media.

Additionally, we examined predicted metabolite secretion for these simulation scenarios. In the minimal medium, *B. pituitosa*, *C. indologenes*, *C. pusillum*, *H. robiniae*, and *S. maltophilia* each produced one type of organic byproduct (acetaldehyde, 4-hydroxy-benzyl alcohol, or 4-hydroxybenzoate). In the exudate medium, *B. pituitosa*, *H. robiniae*, and *S. maltophilia* each expanded their diversity of metabolite production to two or three compounds, including galactose, succinate, and tartrate. Interestingly, *E. ludwigii* and *P. putida* were not predicted to produce any organic byproducts in either the minimal or exudate media, which may reflect the roles they play in the community, explored in the next section. A full list of secreted organic byproducts is found in Table S4 of Supplemental File S1. It should be noted that these findings do not display the full spectrum of each species’ metabolic production potential in the given media, as multiple steady-state metabolic routes can lead to the same objective value (17, 32), and altering nutrient uptake constraints may also influence overflow metabolite production (6).

### Exudate medium supports a more diverse community structure than minimal medium

To assess the potential impact of community coexistence on individual microbial metabolism, as well as the impact of growth medium on community structure, individual metabolic networks were merged into a community model with CarveMe (27). Only models incorporating Biolog data were used based on the findings above. FBA optimizing community growth (the combined growth rates of all seven members) was performed with both the minimal and exudate media to compare the influence of environmental context on predicted steady-state community composition (**Figure 4**). In the minimal medium, optimizing community growth predicted dominance by *B. pituitosa* and *S. maltophilia*, with negligible (several orders of magnitude lower) proportions of *E. ludwigii*, *C. indologenes*, and *H. robiniae*, and zero growth for *C. pusillum* and *P. putida* present. Optimization of community growth on the exudate medium, however, predicted a more diverse distribution of community members, dominated by *B. pituitosa*, *E. ludwigii*, and *S. maltophilia* while also maintaining smaller fractions of *C. indologenes*, *C. pusillum*, and *H. robiniae* (with zero *P. putida*). Experimental measurements have shown *P. putida* to have low abundance in the community (12), though not zero. A potential explanation for the lack of growth observed in the simulation may be limitations in metabolic exchange; *P. putida* was not predicted to produce any organic byproducts in either minimal or exudate medium conditions (Table S4 in Supplemental File S1), and thus its metabolic network may not contribute to the optimization of overall community growth, which is the simulation objective. Compared to monoculture simulations, community simulation with the exudate medium predicted a greater variety of net secreted compounds, including amino acids alanine and leucine, organic acids formate and succinate, and nucleobases thymine and uracil, as well as 4-hydroxy-benzyl-alcohol, 4-hydroxybenzoate, benzaldehyde, and formamide. Predicted net metabolite secretion in the minimal medium was much more limited, including the amino acid leucine and organic acids formate and succinate, as well as 4-hydroxybenzoate.

**Figure 4.**
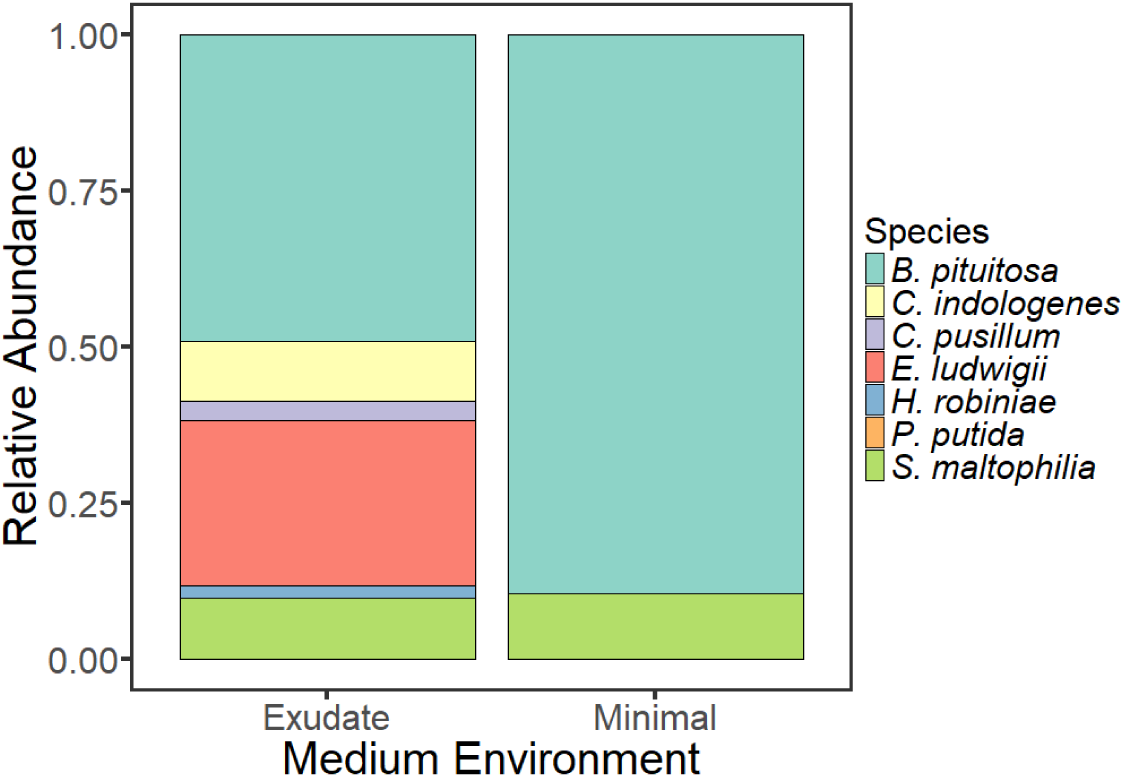
Relative proportions of the seven bacterial members in FBA community model simulations optimizing for total community biomass, performed in either minimal or exudate media. All substrate uptake constraints were set at −10 mmol·gCDW^−1^ ·h^−1^ to examine interactions in a nutrient-rich environment. Species proportions were determined relative to the community growth rate. Note that *C. indologenes*, *E. ludwigii*, and *H. robiniae* were present in minimal medium but at abundances that were orders of magnitude lower than *B. pituitosa* and *S. maltophilia*.

The distinct difference in steady-state composition between minimal and exudate media highlights the important role of nutrient availability in structuring community interactions and also illuminates differences between pure culture and community growth. For example, *E. ludwigii* showed the highest predicted monoculture growth rate of all seven species in the exudate medium (**Figure 3**) but had a lower relative growth rate in the community scenario, which optimized for total community biomass, and was not the most abundant member (**Figure 4**) due to competitive interactions among other members. In the minimal medium (**Figure 5A**), the dominant community members (*B. pituitosa* and *S. maltophilia*) consumed glucose and most of the amino acids, producing relatively few organic byproducts; however, *E. ludwigii* also consumed glucose and malate along with two amino acids and produced several byproducts, some of which were cross-fed to the other community members (ethanol, ornithine, tyrosine, and valine).

**Figure 5.**
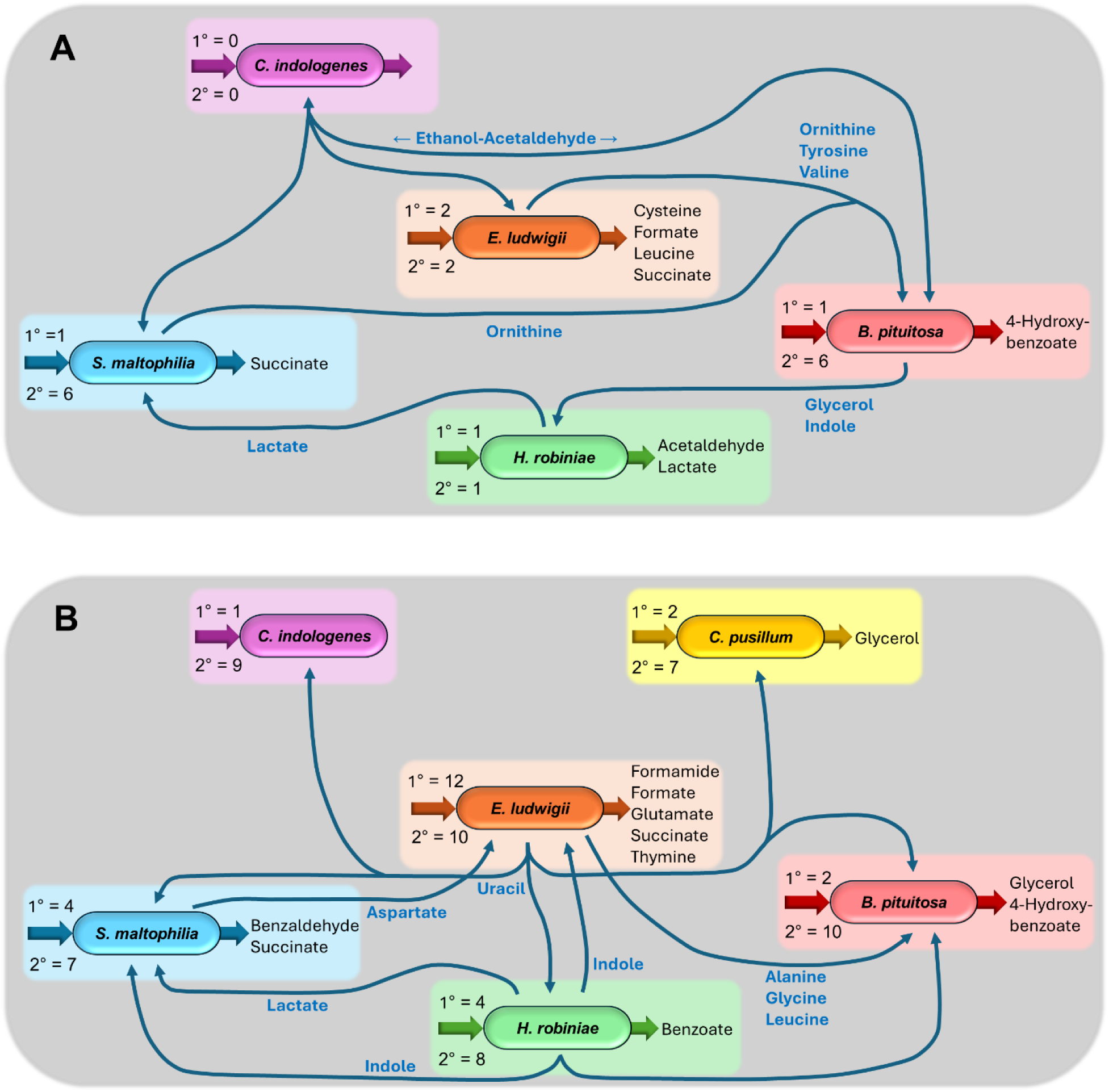
Visual representation of major community dynamics predicted in **(A)** the minimal medium and **(B)** the exudate medium. Intake arrows for each bacterium indicate the number of primary (1°) and secondary (2°) substrates consumed by the species; corresponding output arrows represent compounds that are secreted into the community environment, while blue arrows indicate metabolites exchanged between species (produced by one species and consumed by another). *C. pusillum* and *P. putida* are not pictured in the minimal medium scenario, nor is *P. putida* in the exudate scenario, as they showed zero growth in the community simulation.

The metabolic capacity of each community member increased in the exudate medium compared to the minimal medium, as demonstrated by the increased diversity in byproduct secretion and metabolite exchanges (**Figure 5B**). In the exudate medium, *E. ludwigii* acted as a generalist, consuming the greatest variety of substrates available in the medium: seven sugars (fructose, glucose, lactose, maltose, melibiose, sucrose, and xylose); four organic acids (citrate, fumarate, malate, and oxaloacetate); one alcohol (glycerol); eight amino acids (arginine, asparagine, aspartate, GABA, histidine, lysine, threonine, and tyrosine); and the two nucleosides cytidine and thymidine. Other species consumed a lesser number of substrates, specializing in a small variety of sugars, organic acids, and/or alcohols (arabinose, benzoate, citrate, glycerol, inositol, oxaloacetate, maltose, and ribitol), many of them different from the primary substrates consumed by *E. ludwigii*; their growth was also supported by several amino acids and/or nucleosides. A number of metabolite exchanges including organic acids, amino acids, and nucleobases were also predicted among species (**Figure 5B**), primarily involving *E. ludwigii*, *S. maltophilia*, *H. robiniae*, and *B. pituitosa*. Interestingly, indole was cross-fed from *H. robiniae* to *E. ludwigii*, *B. pituitosa*, and *S. maltophilia*.

### Community structure predicted via exudate medium aligns with *in planta* experimental data

Experiments performed by Niu et al. (12) measured SynCom community member abundance over 13 days. We compared the steady-state predicted simulation abundances (for both minimal and exudate media) with the experimental abundances measured at each time point; we observed an increasing agreement in community member proportions as time progressed, as assessed by Euclidean distance between theoretical and experimental community abundance fractions (**Figure 6**). Data alignment for both minimal and exudate media simulations showed marked improvement toward the end of the experiment as the community matured and stabilized. Distance metrics were substantially reduced for the exudate compared to minimal medium predictions, indicating that the community member abundances predicted by the exudate medium simulation were more similar to the experimental data than were those predicted by the minimal medium simulation. Correlation tests comparing predicted and experimental abundances resulted in higher Pearson’s *r* values for the exudate medium than the minimal medium (*r* = 0.98, 0.98, and 0.97 for the last three time points shown in **Figure 6** and *r* = 0.84, 0.84, and 0.75, respectively), further supporting a robust relationship between the exudate medium predictions and experimental data. These results all together indicate that the root exudate medium enables a more accurate representation of *in planta* community structure as compared to the minimal medium.

**Figure 6.**
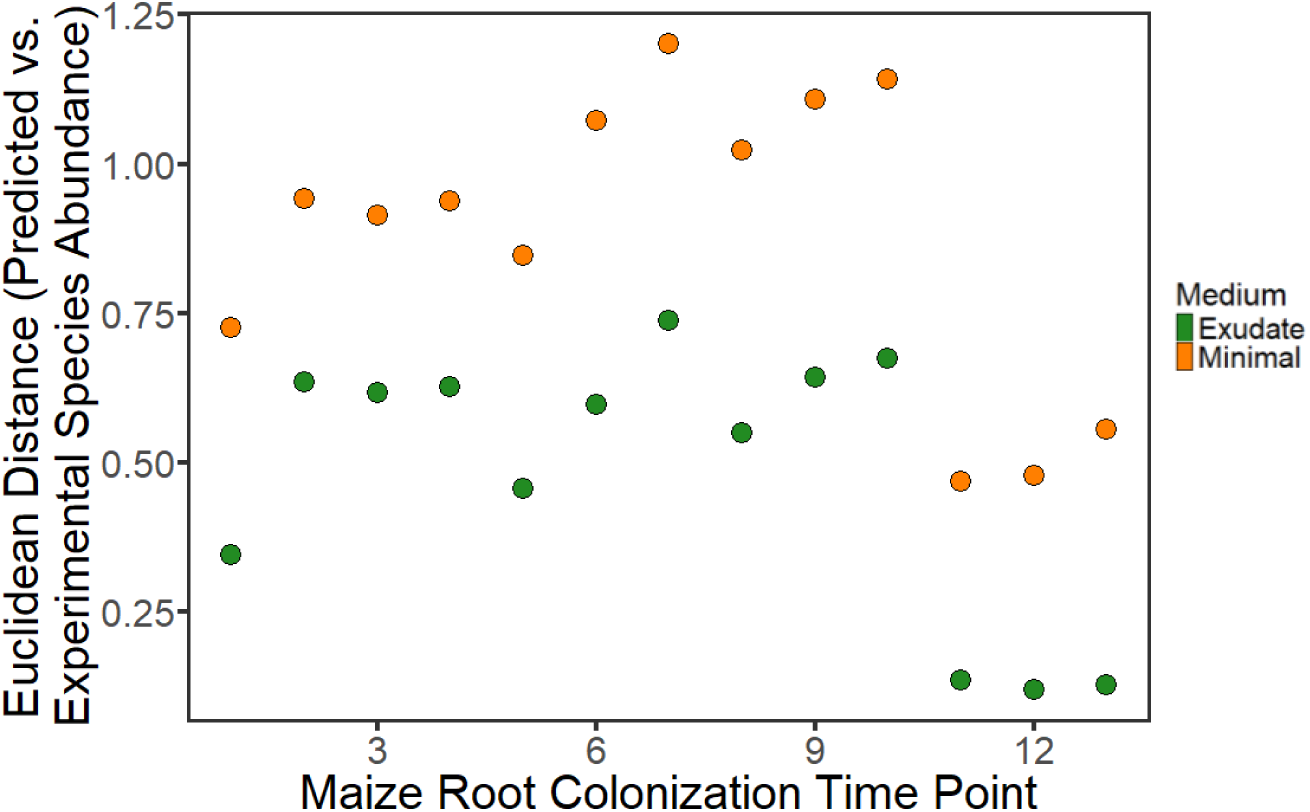
Euclidean distance between experimental and predicted species abundances for community FBA simulations using either minimal or exudate medium (orange and green series, respectively). Simulations were optimized for steady-state community growth to provide predicted relative species abundances; the same simulation result was compared with data from each experimental time point to assess changes in fit across time. Experimental species abundance data, measuring colonization on maize roots over time, was taken from Supplemental Dataset 11 in Niu et al. (12).

## DISCUSSION

This study presents new insights into the biotic and abiotic metabolic interactions in a seven-member bacterial community previously isolated from maize roots (12). To investigate the community-level interactions emerging from individual metabolisms, we reconstructed genome-scale metabolic models for each species and investigated model performance with respect to incorporation of Biolog phenotyping data and the use of different media. Comparing a minimal medium (24) with a custom-designed root exudate medium imitating the plant host environment quantified the effect of environment on simulation output, highlighting the importance of contextual relevance in designing simulation studies. Community FBA was also employed to investigate the impact of the medium type on community structure and predict metabolic exchange between community members in the root environment, providing hypotheses for future experimental testing.

Experimentally, Biolog phenotyping assays showed varied substrate uptake profiles for the seven bacterial species (**Table 2**). When incorporated into metabolic models, these diversified metabolic capacities affected the outcome of FBA predictions (**Figure 2**), with a stronger influence on predictions in the complex root exudate environment than the simpler minimal medium due to the increased number of substrate uptake possibilities. Including experimental data prevents the model construction process from automatically gapfilling the network for all substrates and can help identify genome annotation gaps, reflecting a more accurate physiology and thus improving the hypothesis postulated by the model. Because we cannot predict *a priori* the ways in which experimental data will impact model predictions, collecting data under relevant experimental conditions for a given study is of particular importance.

Medium design is another key aspect of environmental context that requires special attention, as evidenced by the differences between minimal and exudate media simulations observed across species (**Figure 3**). A modeling study focusing on the *Arabidopsis* rhizosphere (33) approached this research question by adding artificial root exudate compounds (consisting of three different sugars, three different organic acids, and three different amino acids (34)) to an existing base media and examining the number of metabolites able to be produced by each bacterial model, finding that the addition of exudate compounds increased the metabolic capabilities of individual species, similar to our results (**Figure 5**, Table S4 in Supplemental File S1). Another prior study investigating apple rhizosphere microbial community interactions *in silico* used metabolomics data to design a medium imitating apple root exudates (35), with FBA-optimized growth rates for a given bacterial species showing inconsistency across simulations with and without exudates, again similar to the patterns observed here.

Along with medium design, specification of nutrient uptake boundaries to reflect the maximum availability of different substrates can also influence simulation results, as demonstrated with the differing number and proportion of substrates when directly comparing the minimal and exudate media (**Figure 3**). The diversity of substrate availability in the exudate medium reduced overall metabolic efficiency by operating multiple catabolic pathways, contributing to the lower growth rates observed in the exudate condition for most species. The *in silico* exudate environment thus more realistically reflects the natural complexity that microorganisms are likely to encounter in the root environment (*i.e.*, smaller amounts of a greater variety of compounds). However, there are other biological complexities the model is unable to capture, such as the effect of dynamic substrate concentrations in a fluctuating environment, transporter kinetics and diffusion limitations on substrate uptake, and gene regulation due to catabolite repression or substrate co-metabolism, which can influence metabolic pathway usage (36, 37). Substrate uptake constraints are often set somewhat arbitrarily if experimental uptake rates are not available; here, we set our maximum overall uptake rate according to the convention of −10 mmol·gCDW^−1^·h^−1^ for the purpose of comparing media performance. This value also lies below the physiologically realistic glucose uptake rate of −18.5 mmol·gCDW^−1^·h^−1^ used in the classic *Escherichia coli* FBA tutorial by Orth et al. (32). It should be noted, though, that in the actual rhizosphere environment, availability of multiple substrates in high concentrations could increase the total substrate uptake rate.

Explicit consideration of the agricultural context through design of a root exudate medium altered predictions for the seven-member community as well as the individual bacterial models. The type of medium affected steady-state community composition, as demonstrated by the ability of the root exudate medium to support a more diverse array of community members than the minimal medium (**Figure 4**). Comparison of species abundances between FBA predictions and data from root colonization experiments (12) exhibited remarkable agreement for the exudate medium simulation, particularly with data from later time points in the experiment when the bacterial community interactions were presumably stabilizing (**Figure 6**). Additionally, examining metabolite exchange (**Figure 5**) showed increased connectivity in the exudate medium with *E. ludwigii* as a metabolic hub, aligning with previous report of its keystone species status (12). Exchange of multiple amino and organic acids and the nucleobase uracil was predicted in the exudate medium, as well as cross-feeding of indole among multiple species.

Indole compounds are known to play important signaling roles within plant hosts (38); in particular, indole is a precursor to the plant hormone indole-3-acetic acid, which is also known to be produced by plant-associated bacteria (39). Interestingly, experimental research applying a consortium of *Bacillus* strains showed a positive effect on walnut seedling growth that was attributed to synthesis of indole-3-acetic acid; this effect was shown to be stimulated via interspecies interactions and was not observed in single-strain experimental controls (40).

While the seven-member SynCom greatly reduces the complexity of the natural community, microbial interactions remain highly complex, and applying genome-scale metabolic modeling within a community context entails inherent limitations. Each individual metabolic model represents a hypothesis regarding a bacterial species’ metabolic capabilities, which is limited by information available in genome annotation databases (41); similarly, the community model provides a hypothesis about how species interact together. It is also important to note that FBA imposes a hypothesis about the model objective which constrains the predicted metabolic fluxes (42), and the solution provided by FBA optimization represents a single possible pathway producing the optimal value for the objective whereas there are typically multiple possible steady-state pathways that can produce the same objective value (and thus the solution represents one prediction of several) (41, 43). Here, we examined the hypothesis of maximal individual or community biomass production (growth rate) under two different nutrient regimes, comparing a typical laboratory environment with a native root exudate scenario. The results we present are therefore one of many possible computational queries, highlighting the flexibility of an *in silico* approach and the opportunities for future work, both in terms of continued computational investigation and experimental validation.

Despite current limitations, the seven-member synthetic community used in this study provides a platform for probing interspecies interactions at the metabolic level to contribute to a more mechanistic understanding of how microbial communities operate. Our findings demonstrate a distinct difference in predicted microbial behavior at both the individual and community levels depending both on the medium environment and on the inclusion of experimental physiological data. Predictions of community composition using a customized root exudate medium, with an expanded suite of substrates compared to a minimal laboratory medium, were better aligned with experimental results and showed expanded metabolite exchange capabilities. The models and simulations presented herein highlight the importance of carefully designing computational investigations to consider the real-world context and research question.

Employing a combination of experimental and theoretical approaches enhances our understanding of the metabolic factors driving host-microbe interactions, which is paramount in further advancing agricultural productivity and engineering more effective conservation and sustainability efforts.

## MATERIALS AND METHODS

### Biolog data collection

Bacterial phenotype data was collected using the Gen III MicroPlate (Biolog Cat. No. 1030) protocol A outlined by Biolog (Biolog, Inc., U.S. Patent # 5,627,045). Briefly, bacterial isolates (*Brucella pituitosa* AA2, DSM 114565; *Chryseobacterium indologenes* AA5, DSM 114485; *Curtobacterium pusillum* AA3, DSM 114566; *Enterobacter ludwigii* AA4, DSM 114484; *Herbaspirillum robiniae* AA6, DSM 114508; *Pseudomonas putida* AA7, DSM 114486; *Stenotrophomonas maltophilia* AA1, DSM 114483) were streaked out on 0.1x tryptic soy agar (TSA) plates and incubated at 30°C for 48 hours. We collected cells from a 3-mm diameter area on the plates using a sterile cotton swab and inoculated Inoculating Fluid A (IF-A; Biolog Cat. No. 72401). We then used a turbidimeter (transmittance set to 100%) to measure turbidity of the inoculated IF-A, with a target turbidity of 90-98%T. We inoculated 100 μL of the inoculated IF-A into each well of the Gen III MicroPlate. Plates were incubated in an OmniLog reader at 30°C for 36 hours and were read at 590 nm every 15 minutes. Biological triplicate data was collected, with replicate averages reported for each species in Supplemental File S2.

### Metabolic model construction

Genome-scale models were constructed for each of the seven bacteria comprising the maize root-associated community developed in Niu et al. (12) based on the genome accessions reported in Niu and Kolter (CP018756, CP018779-86, CP018845-6; the same strains employed in the laboratory work above) (44) using the appropriate Gram-positive or Gram-negative templates in CarveMe (27). Different model versions were constructed for each species, gapfilled and initialized with either a minimal laboratory medium containing glucose and malate as main carbon sources (referred to as minimal medium) or an expanded medium designed to imitate maize root exudates (referred to as exudate medium); gapfilling adds reactions to complete metabolic pathways necessary for growth, and initialization sets basic uptake constraints (27). The minimal medium contained all compounds necessary for growth of individual strains in the laboratory, including several amino acids and vitamins, and was developed specifically to support growth of individual members of the maize root-associated SynCom (24). The compounds included in the exudate medium were identified from literature (26); while not a comprehensive compilation of every known exudate (as the models would be prohibitively complex), representative compounds from a variety of classes were included (sugars, organic and amino acids, alcohols, and nucleosides). Composition of major substrates in both media is reported in **Table 1**, and a comprehensive list of media components is included in Table S1-2 of Supplemental File S1.

Biolog phenotype data were incorporated during the model construction process using the soft constraints option in CarveMe to specify the ability of a microorganism to consume particular compounds (27). Reconstruction employing soft constraints weights corresponding reactions in alignment with phenotyping data, giving preference to substrates the microbe is known to consume experimentally and excluding substrates it is unable to consume. Two model versions were generated for each species: one including phenotyping data through soft constraints and one without data incorporated. The combination of medium type and experimental data inclusion led to four parallel model versions for each of the seven bacterial species: minimal medium incorporating phenotype data, minimal medium without phenotype data, exudate medium incorporating phenotype data, and exudate medium without phenotype data (illustrated in **Figure 1**). Community models were generated with the merge_community command in CarveMe using the previously constructed individual models with incorporation of Biolog data (generating separate community models for minimal and exudate media). The commands used for model generation, as well as the SBML-formatted models themselves, are provided in Supplemental Files S3-4.

### Constraint-based analysis of individual and community models

Flux balance analysis (FBA) simulations were performed in COBRApy with Python 3.9 (45). Models were initialized with carbon substrate uptake bounds set at −10 mmol·gCDW^−1^·h^−1^ to represent replete availability of all carbon substrates. For simulations comparing individual model performance in minimal and exudate media, boundaries were adjusted for a combined maximum uptake of major carbon sources (sugars, alcohols, and organic acids) of −10 mmol·gCDW^−1^·h^−1^, divided equally among the total number of substrates; similarly, the combined maximum uptake of secondary substrates (amino acids, nucleosides, and vitamins) was set to −0.25 mmol·gCDW^−1^·h^−1^, divided equally among the total number of compounds. Individual bacterial model simulations employed the species’ biomass production reaction as the objective function, and community model simulations optimized the community biomass reaction. Supplemental File S3 contains the code used to perform the model simulations reported in this paper.

Resulting flux predictions were exported to Excel for further analysis. Growth rates were compared for individual bacterial models with and without incorporated Biolog data by normalizing to the growth rate predicted without data inclusion: 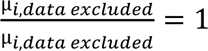 and 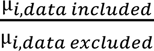 for each pair of simulations, where µ is the simulated growth rate and 𝑖 represents the species. Metabolite consumption and production were examined manually via sorting and identification of exchange reactions among the predicted fluxes (reaction name tags “ex_” in individual models and “ex_”, “abc”, “pts”, “t_”, “t2”, “t5”, and “t6” in community models) to identify metabolite production capabilities and exchange interactions. Relative community member abundance was calculated by dividing each individual species growth rate by the community growth rate. Experimental data regarding relative community member abundance was obtained from Niu et al. (12) (Supplemental Dataset 11), containing species abundance data collected at several different time points during a maize root colonization experiment performed over 13 days. The differences between predicted and experimental community abundances were calculated via Euclidean distance 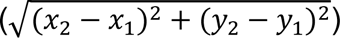 generalized for seven data points, and Pearson’s *r* correlation was used to assess the strength of association between predicted and experimental community abundances.

## ACKNOWLEDGMENTS

This work is supported by Agriculture and Food Research Initiative Agricultural Microbiomes grant no. 2021-67013-34537 from the USDA National Institute of Food and Agriculture and the Novo Nordisk Foundation INTERACT project under grant number NNF19SA0059360.

CReDIT Author Contributions Statement: Conceptualization: AEB, MK; Data curation: AEB, HP, AKG; Formal analysis: AEB; Funding acquisition: AEB, MK; Investigation: AEB, HP, AKG; Methodology: AEB, HP, AKG; Project administration: AEB, MK; Resources: AEB, MK; Software: AEB; Supervision: AEB, MK; Validation: AEB; Visualization: AEB; Writing – original draft: AEB; Writing – review & editing: AEB, AKG, MK

