## Supplemental File S1 for "Genome-scale community metabolic modeling of maize root-associated microbiota shows that root exudates stimulate diverse metabolic interactions"

**Table S1.** Full metabolite composition list for the minimal medium, with compounds arranged in alphabetical order by BiGG identifier. The medium was developed experimentally in Garrell et al. 2026, *Phytobiomes* to enable growth of each of the seven SynCom species in the laboratory. A TSV file is also provided with the supplemental files for direct use with CarveMe.

| BiGG Identifer | Metabolite |
|---|---|
| arg__L | L-Arginine |
| asn__L | L-Asparagine |
| btn | biotin |
| ca2 | Ca2+ |
| cbl1 | Cob(I)alamin |
| cl | Cl- |
| cobalt2 | Co2+ |
| cu2 | Cu2+ |
| CYS | L-Cysteine |
| fe2 | Fe2+ |
| glc__D | D-Glucose |
| gln__L | L-Glutamine |
| h | H+ |
| h2o | H2O |
| ile__L | L-Isoleucine |
| iodine | Iodine c |
| k | K+ |
| leu__L | L-Leucine |
| lys__L | L-Lysine |
| mal__L | L-Malate |
| mg2 | Mg |
| mn2 | Mn2+ |
| mobd | Molybdate |
| na1 | Na+ |
| nac | Nicotinate |
| nh4 | Ammonium |
| no3 | Nitrate |
| o2 | O2 |
| pi | Phosphate |
| pnto__R | (R)-Pantothenate |
| pro__L | L-Proline |
| pydxn | Pyridoxine |
| ribflv | Riboflavin |
| ser__L | L-Serine |
| so4 | Sulfate |
| zn2 | Zn2+ |

**Table S2.** Full metabolite composition list for the root exudate medium, with compounds arranged in alphabetical order by BiGG identifier. The medium was adapted from Hao et al. 2022, *Soil Biology and Biochemistry* to more realistically represent the maize root environment. A TSV file is also with the supplemental files provided for direct use with CarveMe.

| **BiGG Identifier** | **Metabolite** |
|---|---|
| 4abut | 4-Aminobutanoate |
| acon | Cis-Aconitate |
| ala__L | L-Alanine |
| arab__L | L-Arabinose |
| arg__L | L-Arginine |
| asn__L | L-Asparagine |
| asp__L | L-Aspartate |
| bz | Benzoate |
| ca2 | Ca2+ |
| cit | Citrate |
| cl | Cl- |
| cobalt2 | Co2+ |
| cu2 | Cu2+ |
| cytd | Cytidine |
| erthrs | D-erythrose |
| fe2 | Fe2+ |
| fru | D-Fructose |
| fum | Fumarate |
| gal | D-Galactose |
| glc__D | D-Glucose |
| gln__L | L-Glutamine |
| glu__L | L-Glutamate |
| gly | Glycine |
| glyc | Glycerol |
| h | H+ |
| h2o | H2O |
| his__L | L-Histidine |
| ile__L | L-Isoleucine |
| inost | Myo-Inositol |
| iodine | Iodine c |
| isomal | Isomaltose |
| k | K+ |
| lac__L | L-Lactate |
| lcts | Lactose |
| leu__L | L-Leucine |
| lys__L | L-Lysine |

| | |
|---|---|
| mal__L | L-Malate |
| malon | Malonate |
| malt | Maltose |
| melib | Melibiose |
| mg2 | Mg |
| mn2 | Mn2+ |
| mobd | Molybdate |
| na1 | Na+ |
| nh4 | Ammonium |
| no3 | Nitrate |
| o2 | O2 |
| oaa | Oxaloacetate |
| pi | Phosphate |
| rbt | Ribitol |
| ser__L | L-Serine |
| so4 | Sulfate |
| succ | Succinate |
| sucr | Sucrose |
| tartr__M | Meso Tartaric acid C4H4O6 |
| thr__L | L-Threonine |
| thymd | Thymidine |
| tyr__L | L-Tyrosine |
| val__L | L-Valine |
| xyl__D | D-Xylose |
| zn2 | Zn2+ |

**Table S3.** Basic model statistics for each of the model versions constructed: minimal or exudate medium, with or without Biolog phenotyping data. Number of reactions, metabolites, and genes encompassed in the models are reported for each of the seven SynCom species.

| | Number of: | Minimal Medium, excluding Biolog | Minimal Medium, including Biolog | Exudate Medium, excluding Biolog | Exudate Medium, incuding Biolog |
|---|---|---|---|---|---|
| ***B. pituitosa*** | reactions | 2727 | 2542 | 2726 | 2541 |
| | metabolites | 1783 | 1648 | 1783 | 1648 |
| | genes | 1363 | 1332 | 1363 | 1332 |
| ***C. indologenes*** | reactions | 2182 | 2002 | 2181 | 2002 |
| | metabolites | 1460 | 1309 | 1460 | 1309 |
| | genes | 851 | 834 | 851 | 834 |
| ***C. pusillum*** | reactions | 2361 | 2369 | 2360 | 2369 |
| | metabolites | 1575 | 1580 | 1575 | 1580 |
| | genes | 1002 | 1000 | 1002 | 1000 |
| ***E. ludwigii*** | reactions | 2791 | 2794 | 2791 | 2794 |
| | metabolites | 1755 | 1755 | 1755 | 1755 |
| | genes | 1674 | 1674 | 1674 | 1674 |
| ***H. robiniae*** | reactions | 2612 | 2598 | 2609 | 2595 |
| | metabolites | 1684 | 1684 | 1683 | 1683 |
| | genes | 1352 | 1350 | 1352 | 1350 |
| ***P. putida*** | reactions | 2582 | 2809 | 2579 | 2806 |
| | metabolites | 1673 | 1831 | 1673 | 1829 |
| | genes | 1660 | 1691 | 1660 | 1691 |
| ***S. maltophilia*** | reactions | 2530 | 2383 | 2529 | 2382 |
| | metabolites | 1692 | 1557 | 1692 | 1557 |
| | genes | 1008 | 984 | 1008 | 984 |

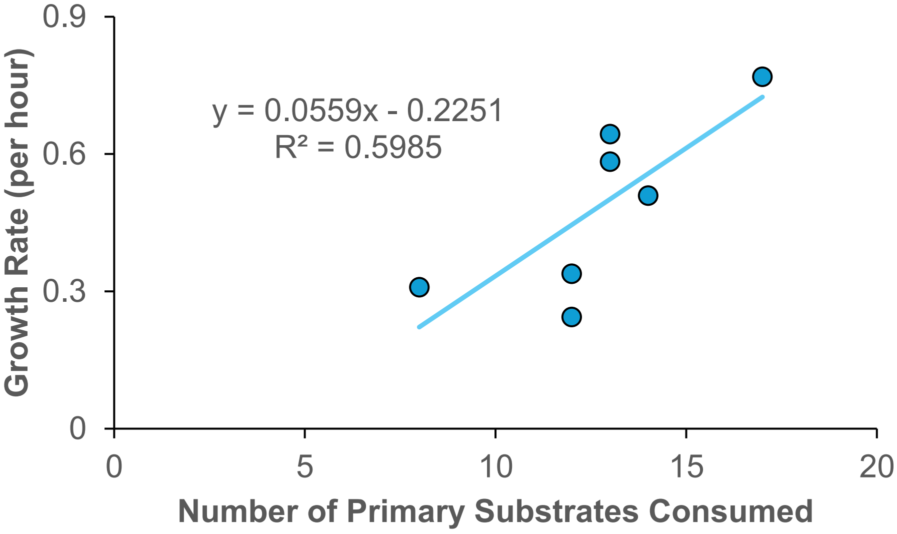


**Figure S1.** FBA-predicted monoculture growth rate of the seven SynCom species in root exudate medium as a function of the number of primary substrates consumed by each species (sugars, alcohols, and organic acids). Linear trendline depicts the line of best fit with an $R^2$ value of 0.60. FBA simulations were performed to maximize growth rate, with the total combined uptake of major carbon sources (sugars, alcohols, and organic acids) set to -10 mmol·gCDW$^{-1}$·h$^{-1}$, and the total combined uptake of secondary substrates (amino acids, nucleosides, and vitamins) set to -0.25 mmol·gCDW$^{-1}$·h$^{-1}$.

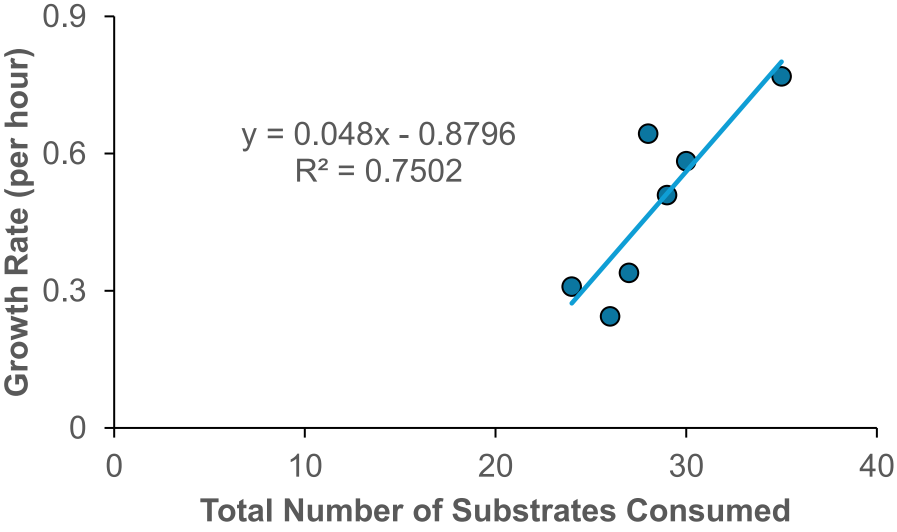


**Figure S2.** FBA-predicted monoculture growth rate of the seven SynCom species in root exudate medium as a function of the total number of primary and secondary substrates consumed by each species (sugars, alcohols, organic and amino acids, and nucleosides). Linear trendline depicts the line of best fit with an $R^2$ value of 0.75. FBA simulations were performed to maximize growth rate, with the total combined uptake of major carbon sources (sugars, alcohols, and organic acids) set to -10 mmol·gCDW$^{-1}$·h$^{-1}$, and the total combined uptake of secondary substrates (amino acids, nucleosides, and vitamins) set to -0.25 mmol·gCDW$^{-1}$·h$^{-1}$.

**Table S4.** Organic byproduct secretion predicted by individual model FBA simulations for each of the seven SynCom species in minimal and exudate media, representing overflow metabolites produced in monoculture. FBA simulations were performed to maximize growth rate, with the total combined uptake of major carbon sources (sugars, alcohols, and organic acids) set to -10 $mmol \cdot gCDW^{-1} \cdot h^{-1}$, and the total combined uptake of secondary substrates (amino acids, nucleosides, and vitamins) set to -0.25 $mmol \cdot gCDW^{-1} \cdot h^{-1}$, to provide a comparable scale for assessing model performance between the two media.

| | Produced in Minimal Medium | Produced in Exudate Medium |
|---|---|---|
| ***B. pituitosa*** | 4-hydroxybenzoate | 4-hydroxybenzoate |
| | | galactose |
| | | tartrate |
| ***C. indologenes*** | acetaldehyde | 4-hydroxybenzoate |
| ***C. pusillum*** | 4-hydroxy-benzyl alcohol | 4-hydroxy-benzyl alcohol |
| ***E. ludwigii*** | - | - |
| ***H. robiniae*** | 4-hydroxy-benzyl alcohol | 4-hydroxy-benzyl alcohol |
| | | galactose |
| ***P. putida*** | - | - |
| ***S. maltophilia*** | 4-hydroxybenzoate | 4-hydroxybenzoate |
| | | succinate |
